# Linking microscopy to diffusion MRI with degenerate biophysical models: an application of the Bayesian EstimatioN of CHange (BENCH) framework

**DOI:** 10.1101/2024.09.30.615704

**Authors:** Daniel Z.L. Kor, Hossein Rafipoor, Istvan N. Huszar, Adele Smart, Greg Daubney, Saad Jbabdi, Michiel Cottaar, Karla L. Miller, Amy F.D. Howard

**Affiliations:** Wellcome Centre for Integrative Neuroimaging, FMRIB, Nuffield Department of Clinical Neurosciences, University of Oxford, Oxford, United Kingdom; Academic Unit of Neuropathology, Nuffield Department of Clinical Neurosciences, University of Oxford, Oxford, UK; Wellcome Centre for Integrative Neuroimaging, Experimental Psychology, Medical Sciences Division, University of Oxford, Oxford, UK; Department of Bioengineering, Imperial College London, London, UK

**Author notes:** Corresponding author: Amy F.D. Howard. Contributed equally (Karla L. Miller, Amy F.D. Howard).

## Abstract

Biophysical modelling of diffusion MRI (dMRI) is used to non-invasively estimate microstructural features of tissue, particularly in the brain. However, meaningful description of tissue requires many unknown parameters, resulting in a model that is often ill-posed. The Bayesian EstimatioN of CHange (BENCH) framework was specifically designed to circumvent parameter fitting for ill-conditioned models when one is simply interested in interpreting signal changes related to some variable of interest. To understand the biological underpinning of some observed change in MR signal between different conditions, BENCH predicts which model parameter, or combination of parameters, best explains the observed change, without having to invert the model. BENCH has been previously used to identify which biophysical parameters could explain group-wise dMRI signal differences (e.g. patients vs. controls); here, we adapt BENCH to interpret dMRI signal changes related to continuous variables. We investigate how parameters from the dMRI standard model of white matter, with an additional sphere compartment to represent glial cell bodies, relate to tissue microstructure quantified from histology. We validate BENCH using synthetic dMRI data from numerical simulations. We then apply it to ex-vivo macaque brain data with dMRI and microscopy metrics of glial density, axonal density, and axonal dispersion in the same brain. We found that (i) increases in myelin density are primarily associated with an increased intra-axonal volume fraction and (ii) changes in the orientation dispersion derived from myelin microscopy are linked to variations in the orientation dispersion index. Finally, we found that the dMRI signal is sensitive to changes in glial cell soma in the WM, but that no parameter in the extended standard model was able to explain this observed signal change, suggesting model inadequacy.

## 1 Introduction

Diffusion MRI (dMRI) provides sensitivity to microstructural tissue features in the brain. However, the interpretation of micrometre-scale features from a signal decay measured in a millimetre-scale MRI voxel is at best challenging, and often ill-posed. To address this, biophysical dMRI models aim to separate measured signals into biologically-meaningful parameters, thereby facilitating investigation into the microstructural underpinnings of dMRI data. These models typically describe the tissue as being composed of multiple compartments. For example, in the brain, white matter models typically aim to describe the intra-axonal space, the extra-axonal space, or cellular soma. The parameters associated with each compartment are intended to relate to biologically meaningful characteristics (e.g. soma radius), and are typically estimated by fitting the model to the dMRI signal..

When modelling microstructure in the long diffusion time regime, the brain white matter (WM) is typically described with the “standard model” [1] as follows. The intra-axonal compartment is modelled as impermeable cylinders of negligible radii (“sticks”) to represent the restricted diffusion of water molecules within axons. The extra-axonal compartment uses an axially symmetric diffusion tensor to capture hindered diffusion in the extracellular space. Both intra- and extra-axonal compartments are typically convolved with a fibre orientation distribution function to model fibre dispersion. If cerebrospinal fluid (CSF) is present, an unrestricted isotropic compartment (“ball”) may also be added, leading to a total of nine free parameters. The WM standard model can be extended to describe other tissue features (e.g. a sphere compartment being used to describe signal contributions from cell soma), though this requires the estimation of more parameters.

Fitting even the standard model to “typical” dMRI data (long diffusion time, multi-shell data with b<4 *ms*/ *μ m*^2^) is very challenging, if not impossible, as the parameter-fitting landscape is often flat [2,3]. Hence, the standard model is degenerate, with multiple parameter sets fitting the signal equally well. One approach to address this issue is to acquire more comprehensive data with an acquisition specifically designed to incorporate useful new information that stabilises the fitting. For example, using multiple diffusion encodings has been shown to provide robust estimates of the standard model parameters [4-6]. For dMRI data that have been acquired using a more “typical” linear diffusion encoding protocol with high angular resolution and/or multiple shells, only relatively few (<5) biophysical parameters can be reliably fitted [7]. Here, a common approach is to constrain parameters such that a restricted biophysical model can be fitted to the dMRI data reliably. Neurite orientation dispersion and density imaging (NODDI) is a constrained form of the standard model, where diffusivities are related and/or fixed to pre-defined values [8]. By adopting these assumptions, NODDI only requires the estimation of five free parameters, facilitating more robust parameter estimation.

These fitted model parameters can then be compared between two conditions (e.g. patients versus controls). However, if the model assumptions are inaccurate or a bad representation of the tissue, the resulting parameter estimates may be biased [7, 9-11] and the groupwise (patient/controls) comparisons misleading. The fact that NODDI fixes the diffusivities can be challenging for both in vivo and postmortem studies, where diffusivities in the latter are largely unknown and biased parameter estimates may lead to spurious relationships when correlated with microscopy.

The Bayesian EstimatioN of CHange (BENCH) framework aims to identify the biological underpinnings of the observed signal changes without having to undertake the highly problematic step of fitting the complete model to the data [12]. BENCH uses a generative model to simulate how the dMRI signal would change if a single model parameter, or set of parameters, was different between the conditions. That is, BENCH defines “change models” in which the change in one model parameter can then be compared with variability in dMRI signals from real data to infer which model parameter best explains the observed change in signal. Crucially, the tissue model does not need to be inverted to infer which parameters are consistent with the observed change, facilitating the use of otherwise degenerate biophysical models that cannot be explicitly inverted.

In this study, we use BENCH to ask whether the dMRI signal measured from the WM possesses sensitivity to the underlying variability in axons and/or cell bodies. Cell bodies or soma are often excluded from data analysis due to increased model degeneracy when including a soma compartment, as it is challenging to distinguish between restricted and unrestricted isotropic compartments. Nevertheless, the density of soma in WM—attributed predominantly to glia—has relevant clinical implications, particularly with respect to neuroinflammation [13]. BENCH provides a framework in which you can identify which model parameters would be implicated in a given biophysical parameter change, where the latter are explicitly derived from microscopy data in the same tissue as dMRI data. In traditional validation studies, a constrained model would be first fitted to the dMRI data voxel-wise, and the fitted parameter estimates correlated with microscopy-derived metrics acquired in the same tissue sample. Here, parameter degeneracy leads to two issues. First, certain compartments—such as a sphere representing glia soma—would often be excluded from the biophysical model to facilitate fitting. Second, model constraints during fitting can again lead to biassed parameter estimates that may affect comparisons with the microscopy.

Circumventing these issues, we developed “continuous BENCH” to relate changes in dMRI signal to variations in the density and dispersion of axons or cell soma extracted from microscopy. To accomplish this, we first adapt the existing BENCH framework to infer change that corresponds to continuous, rather than categorical, variables derived from microscopy data. Continuous BENCH is first tested on synthetic data generated using numerical simulation of microstructure mesh substrates, providing proof-in-principle of detecting axon and soma density. Continuous BENCH was then applied to an ex-vivo macaque brain with co-registered dMRI and microscopy data. The specific microscopy stains target axons and cell soma, enabling the derivation of metrics of dispersion and density. These metrics (termed “continuous variables”) provided the axis along which we characterise the direction of change in our measured dMRI signal. We then used continuous BENCH to estimate which biophysical model parameter best explains the measured change in the dMRI signal due to microscopy-derived differences in axons and glia.

## 2 Continuous BENCH

The continuous BENCH pipeline consists of three stages (Figure 1): the training stage (green box; “Training”), setting up a generalised linear model (GLM) (beige box; “Continuous change”), and the inference stage (blue box; “Inference”). Note that vectors are given in bold. The hat notation is used to represent a unit change in the given vector (i.e. a direction).

**Figure 1:**
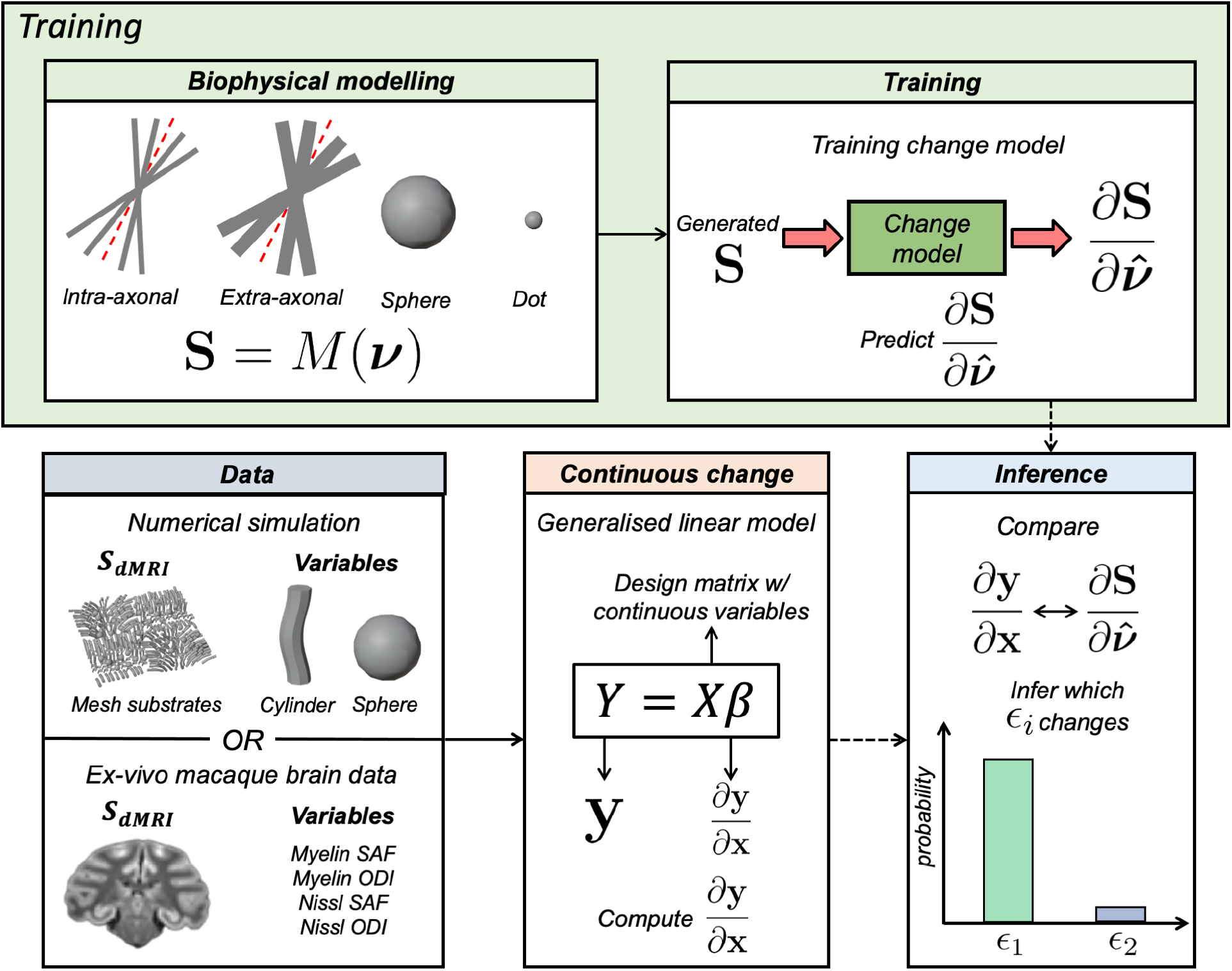
An overview of continuous BENCH. First, in BENCH’s training stage (green box; “Training”), we aim to infer changes in the generated dMRI signal with respect to changes in biophysical parameters 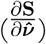. This is achieved by training “change models” (i.e., regression models) for each biophysical parameter (*ϵ*_*i*_) from a designated diffusion biophysical model (*M*) using generated data. Next, we quantify how the measured dMRI signal changes due to our continuous variables-of-interest 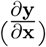 using a generalised linear model (GLM) (beige box; “Continuous change”). In this study, we separately consider numerically simulated and real ex-vivo macaque brain data (blue box; “Data”). The simulated data is to demonstrate continuous BENCH under conditions where the ground truth is known, while the ex-vivo brain data shows continuous BENCH applied to real data. Finally, the inference stage (blue box; “Inference”) uses Bayes’ rule to calculate the probability of a given biophysical parameter driving the measured signal variability.

First, BENCH’s training stage (Section 2.2) uses a generative model (*M*) to train “change models” where, for a set of input biophysical parameters (**ν**), we generate a dMRI signal (**S**) and a series of “change vectors” 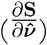 which describe the change in the dMRI signal induced by a unit change 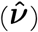 in the biophysical model parameters (***ν***). Next, in real data (**y**), we use a generalised linear model (GLM) (Section 2.3) to compute how the observed dMRI signal changes with respect to some continuous variable(s) of interest (**X**). In our case, **X** is the axon or cell body density derived from microscopy, or both. The output of the GLM 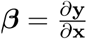 defines the “direction of change” of interest. Finally, BENCH’s inference stage (Section 2.4) compares the observed 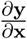 with the predicted outputs from change models 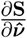 to infer which biophysical parameter (if any) best explains the observed differences in our dMRI signal due to the continuous variables-of-interest. The quality of fit, measured by the chi-squared distance between predicted and observed changes, determines how well the biophysical parameter set 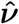 explains the dMRI signal differences.

The crux of BENCH is that by reframing the question to consider the MRI signal change across conditions (across subjects, timepoints or voxels), rather than fitting the model to each datapoint separately, we can circumvent inverting the model. This enables investigation of otherwise intractable models. BENCH (i) assumes that, regardless of the absolute parameter values, a change in a given set of parameters alters the signal in a consistent direction, and (ii) aims to identify candidate parameters consistent with the observed change. If multiple parameters could be responsible, BENCH can only make a statement about the probability of each.

### 2.1 Theory

Given a biophysical model (*M*) that predicts a dMRI signal (**S**) with input biophysical parameters (***ν***), **S** = *M*(***ν***), we aim to identify one set of biophysical parameters whose change 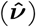 best predicts a measured change in the dMRI signal (**Δy**) relative to a baseline dMRI measurement **y**. Note that **S** represents the model-predicted dMRI signal, while **y** is the measured dMRI signal.

Details of BENCH have been previously described in [12]. In BENCH, we bypass model inversion by instead inferring 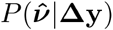, the probability of a change in given biophysical parameter 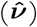 given a baseline dMRI signal (**y**)and a measured change in dMRI signal (**Δy**). This process is formalised using Bayes’ rule:

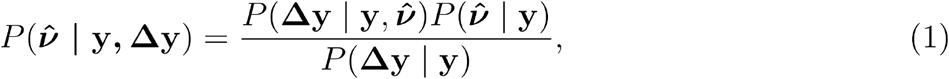

where the denominator is a normalisation constant (often called the evidence), and 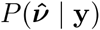 is the prior on the parametric change 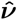. In principle, the prior can be any form of distribution. We assumed a uniform distribution, which implies that no specific change is more likely than any other, given any measured baseline dMRI signal. Hence, we only need to estimate the likelihood 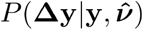. This can be achieved by first computing:

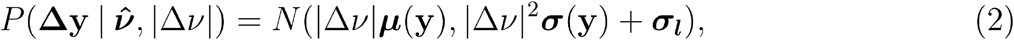

where |Δ*ν*| is the magnitude of change for a change in parameter set ***ν*** of arbitrary size:

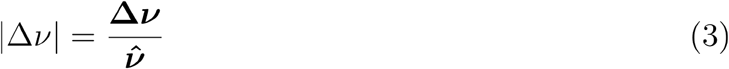

*N*(***μ, σ***) is a normal distribution, with mean ***μ*(y)** and variance ***σ*(y)**, that describes the spread in of possible change vectors originating from some baseline signal **y** (Figure 2). ***σ***_***l***_ is the noise covariance matrix of the continuous variable (indexed *l*) as estimated from the data (Section 4).

**Figure 2:**
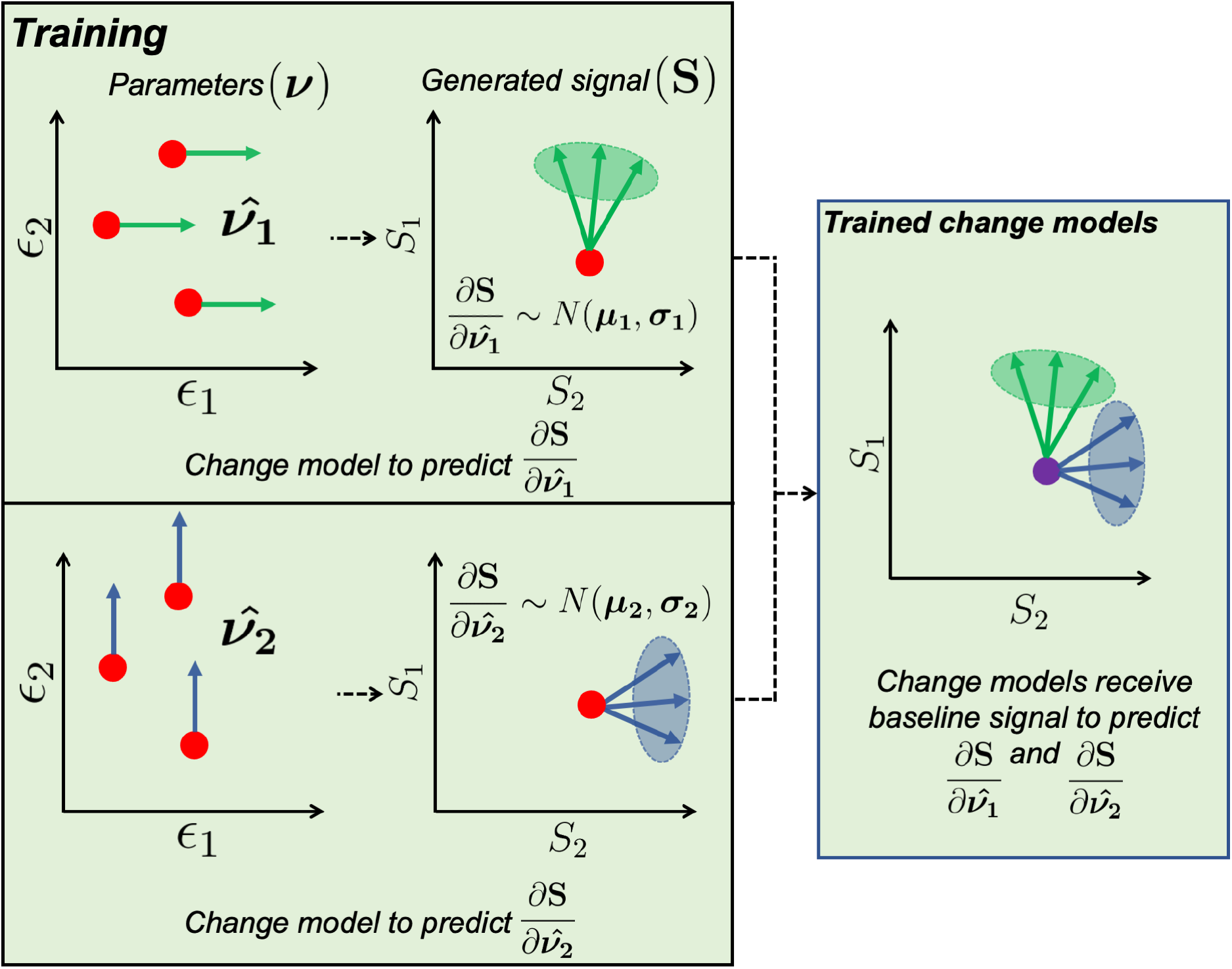
Illustration of the training stage. This is done in the context of a degenerate diffusion biophysical model with biophysical parameters *ϵ*_1_ and *ϵ*_2_. As the model is degenerate, several parameter combinations (red points, parameters graph) generate the same signal (red point, generated signal graph). During the training phase (box titled “Training”), we aim to train a separate change model for each parameter. For example, while training the model for *ϵ*_1_, we calculate the signal for a given pair of *ϵ*_1_ and *ϵ*_2_ and the signal change due to a small perturbation in *ϵ*_1_ only (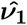, a green arrow in the parameters graph) to produce a change vector in the signal space 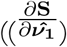 a green arrow in the generated signal graph). If we repeat this for many combinations of *ϵ*_1_ and *ϵ*_2_, we can estimate a normal distribution of change vectors (green ellipse *N*(***μ***_**1**_, ***σ***_**1**_)). This process is repeated for all parameters in our biophysical model (e.g. *ϵ*_2_) to generate their corresponding change models. These trained change models (box titled “Trained change models”) can thus provide robust estimates of which parameter has changed (if the parameter distributions/Gaussians are distinct and point in different directions) even when the baseline signal (purple point) itself is degenerate.

Since we only aim to identify if a parameter is changing, rather than the magnitude of that change, we marginalise 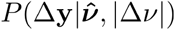 with respect to |Δ*ν*| to get 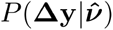. We estimate ***μ*(y)** and ***σ*(y)** using trained change models, as outlined below (Section 2.2). Please refer to [12] for detailed derivations.

### 2.2 Training change models

As described above, BENCH consists of three stages: training, GLM, and inference. The objective of the first (training) stage is to estimate a distribution of change vectors *N*(***μ, σ***) for each biophysical parameter. Figure 2 outlines how this is done in the context of an example degenerate biophysical diffusion model that has input parameters *ϵ*_1_ and *ϵ*_2_ (i.e. ***ν*** *=* [*ϵ*_1_, *ϵ*_2_]) and outputs the dMRI signal(s) **S** = [S_1_, S_2_]. For example, *ϵ*_1_ and *ϵ*_2_ could represent diffusivity and dispersion, while **S** represents the powder average dMRI signal for two different b-values. Because this is a degenerate model, multiple parameter sets (combinations of *ϵ*_1_ and *ϵ*_2_) may lead to the same generated dMRI signal **S**.

In practice, a separate training dataset is generated for each change model (indexed *i*, where *i* = 1, …, *N*) *N* is the total number of change models considered. Typically, *N* corresponds to the number of parameters in the model, but this may not always be the case. For example, *N* can be larger if the models capture interactions where one parameter increases as another decreases.

In this study, we trained individual change models that either characterise a change in a single biophysical parameter or a change in two parameters. The latter case applies only to signal fractions, as the signal fractions are normalised to sum to one. Thus, a change in one compartment’s signal fraction necessitates a complementary change in another compartment’s signal fraction, such as (*f*_*in*_ − *f*_*ex*_), (*f*_*in*_ − *f*_*sph*_), or (*f*_*ex*_ − *f*_*sph*_).

For change model *i*, the training dataset is generated using the following steps:

1. Sample a parameter combination (***ν*** *=* [*ϵ*_1_, *ϵ*_2_, …]) from a set of prior distributions. These distributions are predefined empirically.
2. Generate a baseline dMRI signal (**S** = *M*(***ν***))
3. Induce a small change in the chosen parameter 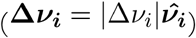. The magnitude of change is drawn from a uniform distribution. Here, **Δ*ν***_***i***_ represents a change in the parameter combination required as training data to train change model *i*.
4. Generate a new dMRI signal (**S** = *M*(***ν*** *+* **Δ*ν***_***i***_)) with the new parameter combination.
5. Compute the corresponding change vector 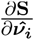, given **Δ*ν***_***i***_, using:

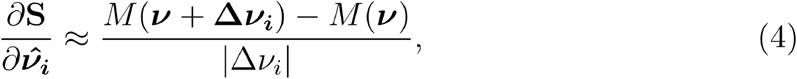

where we assumed that dMRI signals change linearly with the parameter.
6. Repeat steps 1 to 5 to generate a predefined number of pairs of (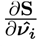, **S**_**a**_).

These steps are repeated for all parameters of the biophysical model *M*.

Up to this point, we have estimated 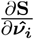 for a discrete set of generated **S**. To describe the distribution of change vectors *N*(***μ***_***i***_, ***σ***_***i***_) for any **S**, we assume that ***μ***_***i***_ and ***σ***_***i***_ will vary smoothly over signal space such that ***μ***_***i***_**(*S*)** and ***σ***_***i***_**(*S*)** can be estimated using regression models trained on our generated pairs (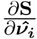, **S**). For ***μ***_***i***_**(*S*)**, the regression model is:

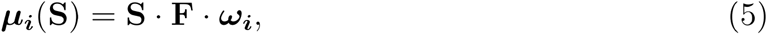

where **S · F** is the design matrix. **F** is a linear transformation of **S**, where **S · F** produces a basis set that can characterise ***μ***_***i***_**(*S*)** varies over **S. *ω***_***i***_ are the regression weights, with dimensions defined by the number of variables in the basis set (rows) and the number of dMRI summary measures (columns).

***σ***_***i***_**(*S*)** is estimated with a similar formulation, albeit with an additional transformation *T* to ensure that ***σ***_***i***_**(*S*)** is a positive definite matrix:

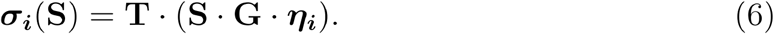

**G** is another linear transformation. In order to account for the positive definite nature of ***σ***_***i***_**(*S*), T** includes the arrangements of elements, exponentiation of the diagonals and the matrix multiplication for the inverse Cholesky decomposition. ***η***_***i***_ are the weights, with the row length set by the number of independent variables and the number of columns corresponding to the unique elements in the covariance matrix ***σ***_***i***_.

Inserting these regression models into the normal distribution *N*(***μ***_***i***_**(*S*), *σ***_***i***_**(*S*)**):

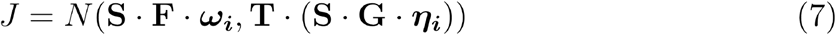

where the likelihood of observing 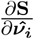 is:

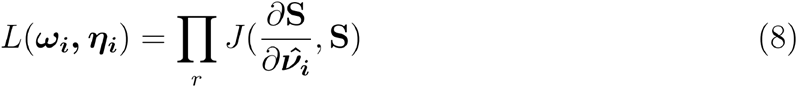

The optimal weights ***ω***_***i***_, ***η***_***i***_ are estimated using a maximum likelihood optimisation and 𝓇 pairs of (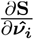, **S**) from the training dataset. These weights can then be used to estimate ***μ***_***i***_ and ***σ***_***i***_ at any observed baseline dMRI signal **y** derived from real data via Equations 5 and 6, respectively.

### 2.3 Extending BENCH to continuous variables

Next, we use real data to define the baseline dMRI signal **y** and signal change **Δy** that are meaningful to our research question (**y** is used to describe the observed diffusion signal, whilst **S** is the generated signal from the forward model). This allows us to calculate the signal change 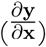 with respect to variable(s)-of-interest **x**.

Previously, Rafipoor et al. defined **x** to be a categorical variable describing two discrete groups (healthy controls and patients) [12]. Specifically, **x** was a vector where healthy controls were denoted as 0 and patients represented by 1. However, many research problems cannot be formulated in terms of categorical variables of this kind. Here, we extend the BENCH framework to consider continuous variables (here, histological stains that may have relevance to dMRI signals). We relate changes in the dMRI signal to continuous variables-of-interest via a general linear model (GLM), in the case where there are *q* continuous variables:

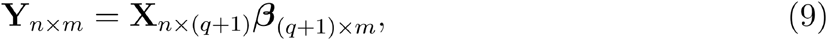

where:

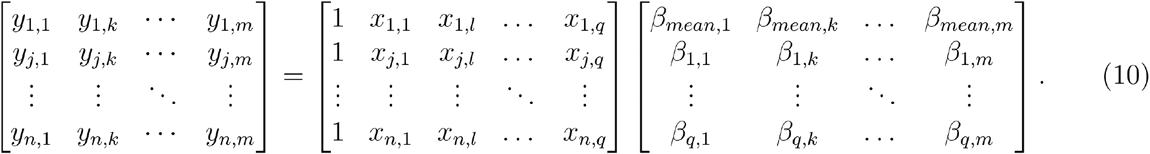

Here, *j* = 1, …, *n* are the different measurements (e.g. subjects or voxels) for which we have *k* = 1, …, *m* dMRI summary measures (as described later in Section 3.1) and *l* = 1, …, *q* continuous variables. As with any GLM, the *β*s are estimated by multiplying the matrix of summary measures, **Y**, with the pseudoinverse of the design matrix, **X**.

### 2.4 Inference

The inference stage (Figure 3) combines outputs from the training stage 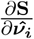 and the GLM 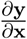 to infer which change model best explains the observed change in dMRI with respect to the continuous variables-of-interest.

**Figure 3:**
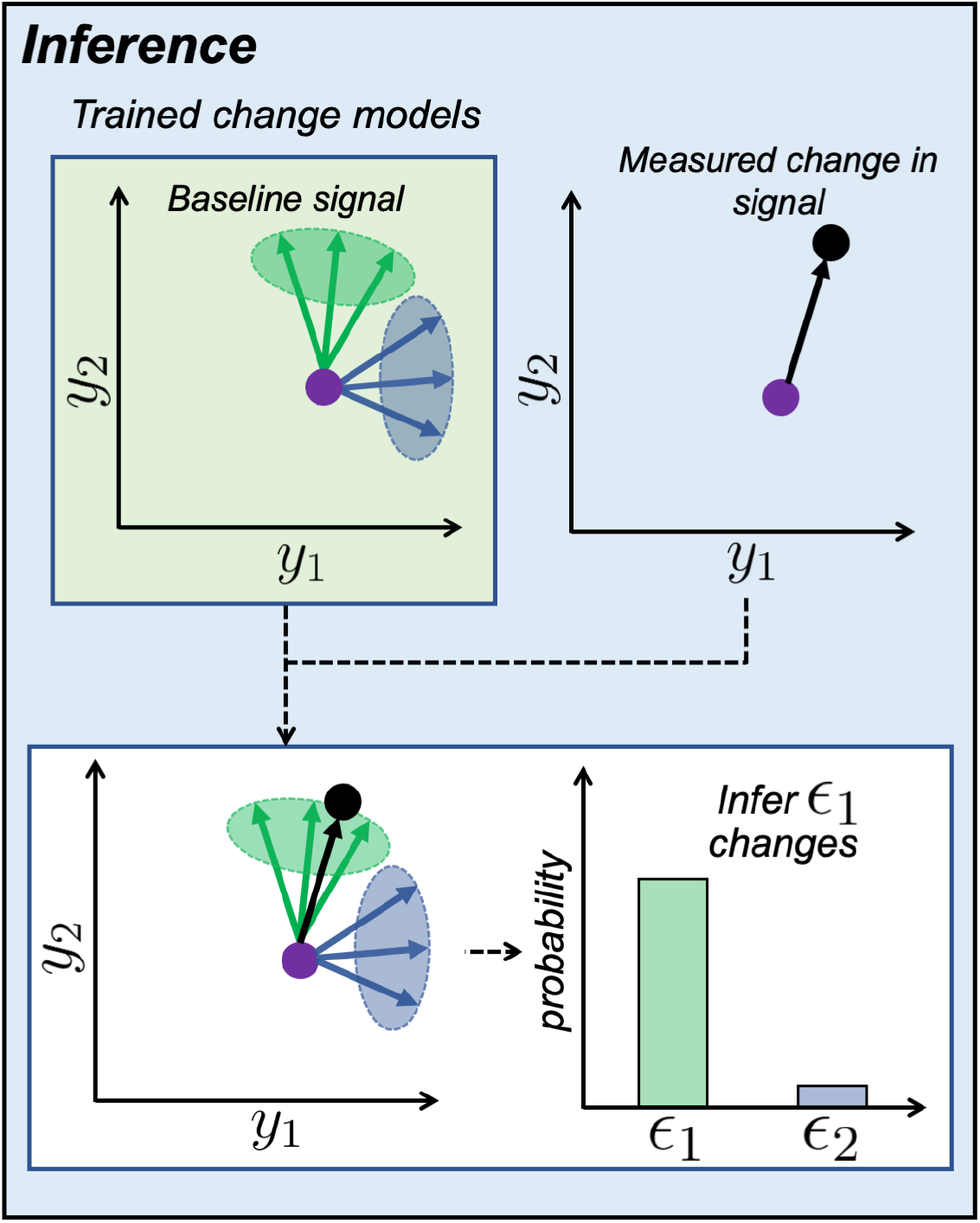
Illustration of the inference stage. With the trained change models and a baseline dMRI signal (purple dot), we can generate a distribution of change vectors for *ϵ*_1_ (green ellipse) and *ϵ*_2_ (blue ellipse). We then compare a measured dMRI signal change associated with a continuous variable (purple to black dot) with the distribution of change vectors. This allows us to estimate the probability of each biophysical parameter explaining the measured change in the dMRI signal (Equation 1). In this case, we would infer that the observed dMRI signal change is mostly likely due to a change in *ϵ*_1_.

The first row of the GLM, ***β***_***mean***_ =[*β*_*mean*,1_ … *β*_*mean,m*_], is our data-driven estimate of the baseline dMRI signal. ***β***_***mean***_ is inputted into the trained change models to determine *N*(***μ***_***i***_(***β***_***mean***_), ***σ***_***i***_(***β***_***mean***_)) for each biophysical parameter.

Next, we compute the measured change in dMRI signal (relative to ***β***_***mean***_) due to each continuous variable *x*_*l*_. This is achieved by multiplying the direction of change 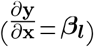 with the change effect size (*δ*_*l*_), where *δ*_*l*_ is measured directly from the data. To account for measurement noise in the dMRI signal, a noise covariance matrix (***σ***_***l***_ in Equation 2) is also estimated directly from the data. The estimation of *δ*_*l*_ and ***σ***_***l***_ is described later in Section 4.

Putting this together (Equations 2), for each continuous variable *l* and change model *i*, we evaluate the likelihood 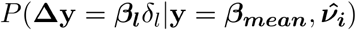. Using this likelihood in Bayes’ rule (Equation 1) gives the probability of a change in biophysical parameter *ϵ*_*i*_ explaining our data, 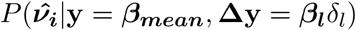. These likelihoods are normalised across all biophysical parameters and inference is performed separately for each continuous variable-of-interest.

## 3 Data simulation and acquisition

### 3.1 Numerical simulations

Monte Carlo simulations [14] of spins diffusing in a geometric space designed to mimic the WM were performed to establish whether our method can detect subtle changes approximating the density of axons and soma. A total of 24 artificial 3D meshes (Figure 4) were created with axons as undulating cylinders (radius = 2.5 *μm* [15]; undulation amplitude = 1 *μm*) and cellular soma as spheres (radius = 1.8 *μm*, which is within the observed range of glia cell soma size [16]). Axons were oriented in each substrate to simulate fibre dispersion, approximately following a Watson distribution [8]. Substrates (size = 200 × 200 × 25 *μm*^3^) were generated for various numbers of undulating cylinders (n=[128, 168, 208]) and spheres (n=[0, 20, 60, 100, 140, 180, 220, 260, 300, 340, 380, 420]), which resulted in substrates with varying intra-axonal (0 to 15% of substrate) and sphere (or intra-soma) volume fractions (0 to 1.5% of substrate, in line with microglia cell density we previously observed in postmortem human brain WM [17]). Simulations were performed using 160,000 spins (uniformly initialised in both intra- and extra-axonal space) with a time step of 8.62 *μs* [18] and impermeable membranes. To simulate ex-vivo tissue conditions, bulk water diffusivity was set to 0.8 *μm*^2^*/ms* dMRI signals were simulated for each substrate using acquisition parameters very similar to what was previously used to acquire ex-vivo macaque brain HARDI data in [19], except with a reduced number of gradient directions: 90 gradient directions acquired each at b = 7 and 10 *ms/μm*^2^ and 18 images acquired at b = 0 *ms/μm*^2^. Other parameters include: Δ = 24 *ms, TE* = 42.5 *ms, G* = 15.9, 19.1 *G*/*cm*, δ = 14 *ms*.

**Figure 4:**
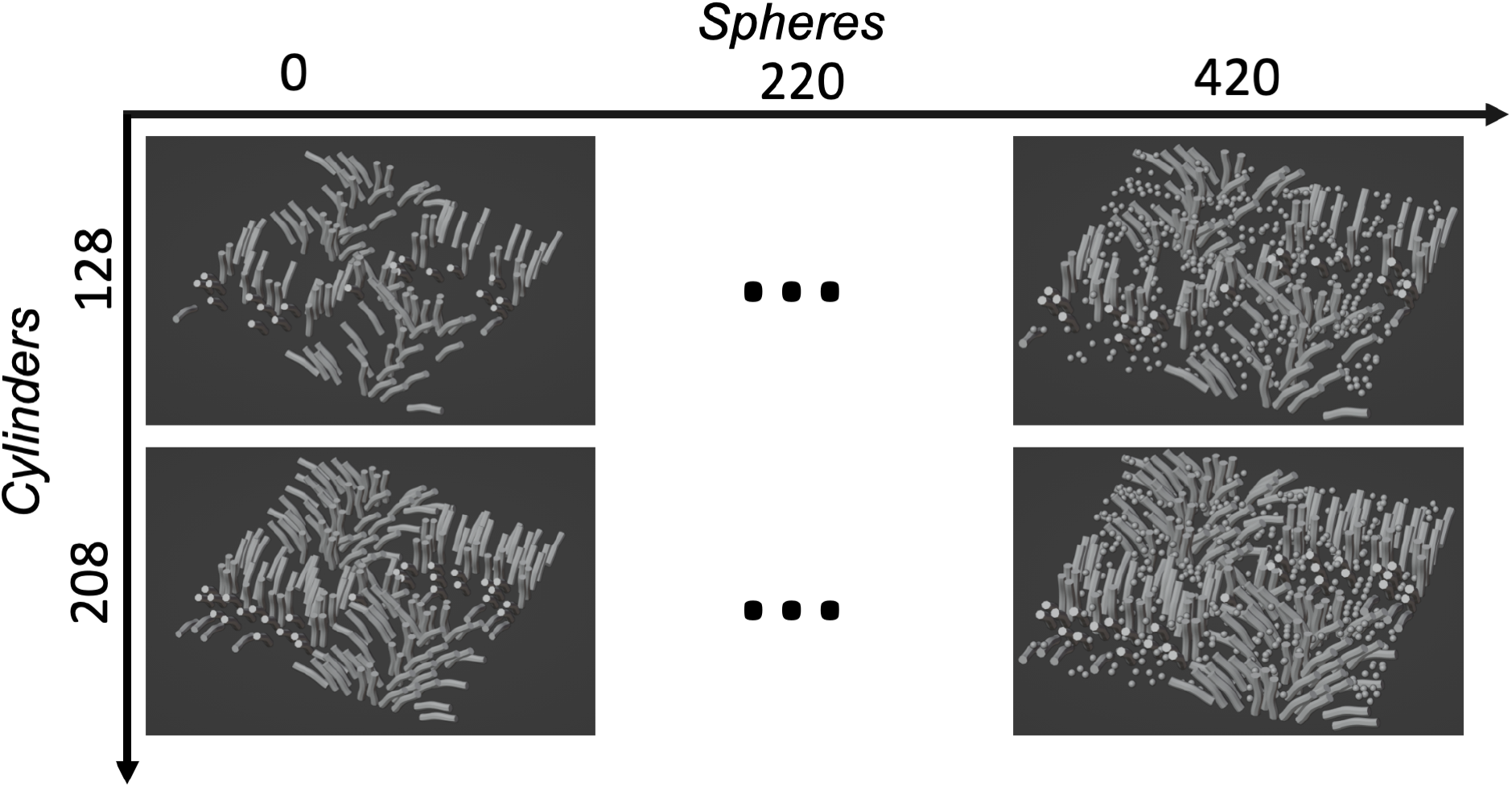
The mesh substrates used for simulation. A total of 24 substrates were used, each with a different number of cylinders (representing myelinated axons; radius = 2.5 *μm* [15]) and spheres (representing glial cell soma; diameter =1.8 *μm* [16]).

BENCH is designed to work with rotationally invariant summary measures of the signal, rather than the raw dMRI signals themselves. To achieve this, spherical harmonics [20, 21, 1] were fitted to the dMRI signal and the summary measures were calculated as the weighted average of the squared coefficients for degree *ℓ* = 0 and 2. This resulted in 4 rotationally invariant summary measures (2 summary measures × 2 shells) per mesh substrate. For more details, please refer to the Appendix.

### 3.2 Ex-vivo macaque brain data

We applied continuous BENCH to previously published ex-vivo macaque brain data [19], where postmortem dMRI and co-registered [22] microscopy metrics specific to cell soma and myelin were available. Note that we only describe parts of the data that are directly relevant to this work.

#### Diffusion MRI data

The acquisition and preprocessing of the ex-vivo HARDI data from a perfusion-fixed brain of an adult rhesus macaque have been previously described in [19]. Briefly, data were acquired on a 7T small animal scanner (Agilent) with 1000 gradient directions at b = 7, 10 *ms/μm*^2^, along with 80 images acquired at negligible diffusion-weighting (b = 0 ms/*μm*^2^). Images were acquired with 1 mm isotropic resolution. Other acquisition parameters are identical to what was used for the numerical simulation. A WM mask was generated [19]. Rotationally invariant summary measures were similarly calculated from the dMRI signal (c.f. Appendix).

#### Microscopy data

Following MRI data acquisition, the macaque brain was sectioned along the anterior-posterior axis to produce coronal tissue sections. Of relevance to this study, 50 *μm* thick sections were histologically stained for Cresyl violet to visualise Nissl bodies, and Gallyas silver, which targets myelin [19]. Each stain was repeated every 350 *μm* throughout the brain. Sections were imaged at a resolution of 0.28 *μm* per pixel. All microscopy slices were co-registered to the MRI volume using TIRL [22].

The histology images were analysed to extract two quantitative metrics: the stained area fraction (SAF) and the orientation dispersion index (ODI). The SAF is a metric for microstructural stain density, while the ODI describes how dispersed the stained microstructure is. The details of how both SAF and ODI are derived are published in [16]. In brief, stain segmentation was performed using data-driven thresholds based on a weighted-Otsu method [23], similar to that implemented in [17]. The SAF was then calculated as the number of positive pixels over a local neighbourhood the size of the MRI voxel. Dispersion was quantified by performing structure tensor analysis to estimate the primary fibre orientation per pixel [24]. A Bingham distribution was then fitted to the set of microscopy-derived orientations within the neighbourhood of a MRI voxel to calculate an ODI that is directly comparable with MRI-derived fibre orientation distributions, albeit in 2D.

Continuous BENCH was applied to co-registered dMRI, Nissl-derived, and myelin-derived data from the anterior brain (17,945 voxels in total). A voxel mask is shown in Appendix Figure S1 (myelin staining in the posterior brain was excluded as it was found to be corrupted).

## 4 Data analysis

We applied continuous BENCH to both numerical simulation and ex-vivo macaque brain data. For either dataset, we need to specify:

1. a generative biophysical model *M* used for training change models (c.f Section 2.2)
2. the measured direction of change ***β***_***l***_ (c.f Section 2.3) and the uncertainty on ***β***_***l***_ (i.e. ***σ***_***l***_, the noise covariance matrix for *l*^*th*^ continuous variable) (c.f Section 2.4)

### 4.1 Biophysical model

For both datasets, we used an extended WM standard model that included an intra-axonal compartment modelled as dispersed sticks, an extra-axonal compartment characterised as dispersed zeppelins, and an intra-soma compartment modelled as a sphere using the Gaussian phase distribution approximation [25-27]. A dot compartment was also added to account for stationary water, which is often present in ex-vivo data [28, 29]. All of these compartments were characterised as either anisotropically (zeppelin, sticks) or isotropically (sphere, dot) restricted diffusing spins. When analysing data simulated from mesh substrates, we also included an isotropic Gaussian (ball) compartment to model freely diffusing spins that do not interact with any membrane. In total, our biophysical models require 10 and 12 model parameters for simulated and real ex-vivo data, respectively (Table 1). In both cases, the signal fractions were assumed to sum to 1.

**Table 1:** Parameters of the extended WM standard model. When analysing simulation data, a ball compartment (parameters shaded in grey) was added to explicitly model spins that do not interact with any membranes.

| Parameter | Units | Description | Range |
| --- | --- | --- | --- |
| $f_{in}$ | - | Signal fraction for intra-axonal compartment | [0, 1] |
| $f_{ex}$ | - | Signal fraction for extra-axonal compartment | [0, 1] |
| $f_{sph}$ | - | Signal fraction for intra-soma compartment | [0, 0.10] |
| $f_{dot}$ | - | Signal fraction for the dot compartment | [0, 0.10] |
| $f_{ball}$ | - | Signal fraction for ball compartment | [0, 0.10] |
| $ODI$ | - | Orientation dispersion index | [0.01, 0.50] |
| $D_{a,in}$ | $\mu m^2/ms$ | Axial diffusivity for the intra-axonal compartment | [0.50, 1.5] |
| $D_{a,ex}$ | $\mu m^2/ms$ | Axial diffusivity for the extra-axonal compartment | [0.50, 1.5] |
| $D_{r,ex}$ | $\mu m^2/ms$ | Radial diffusivity for the intra-axonal compartment | [0.50, 1.5] |
| $D_{sph}$ | $\mu m^2/ms$ | Isotropic diffusivity for the intra-soma compartment | [0.50, 1.5] |
| $D_{ball}$ | $\mu m^2/ms$ | Isotropic diffusivity for the ball compartment | [0.50, 2.5] |
| $R_{sph}$ | $\mu m$ | Radius of the intra-soma compartment | [0.01, 10] |

Added when modelling simulation data

### 4.2 Training change models

Each change model was trained using 20,000 pairs of baseline dMRI signal and generated change vector (**S**, 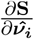) produced from the generative biophysical model (c.f. Section 2.2). We trained a change model for each biophysical parameter, and combinations of signal fractions (e.g. sphere fraction replacing extra-axonal fraction (*f*_*sph*_ − *f*_*ex*_)) such that the signal fractions sum to 1. These change models are referred to and explained in Figure 5 caption.

**Figure 5:**
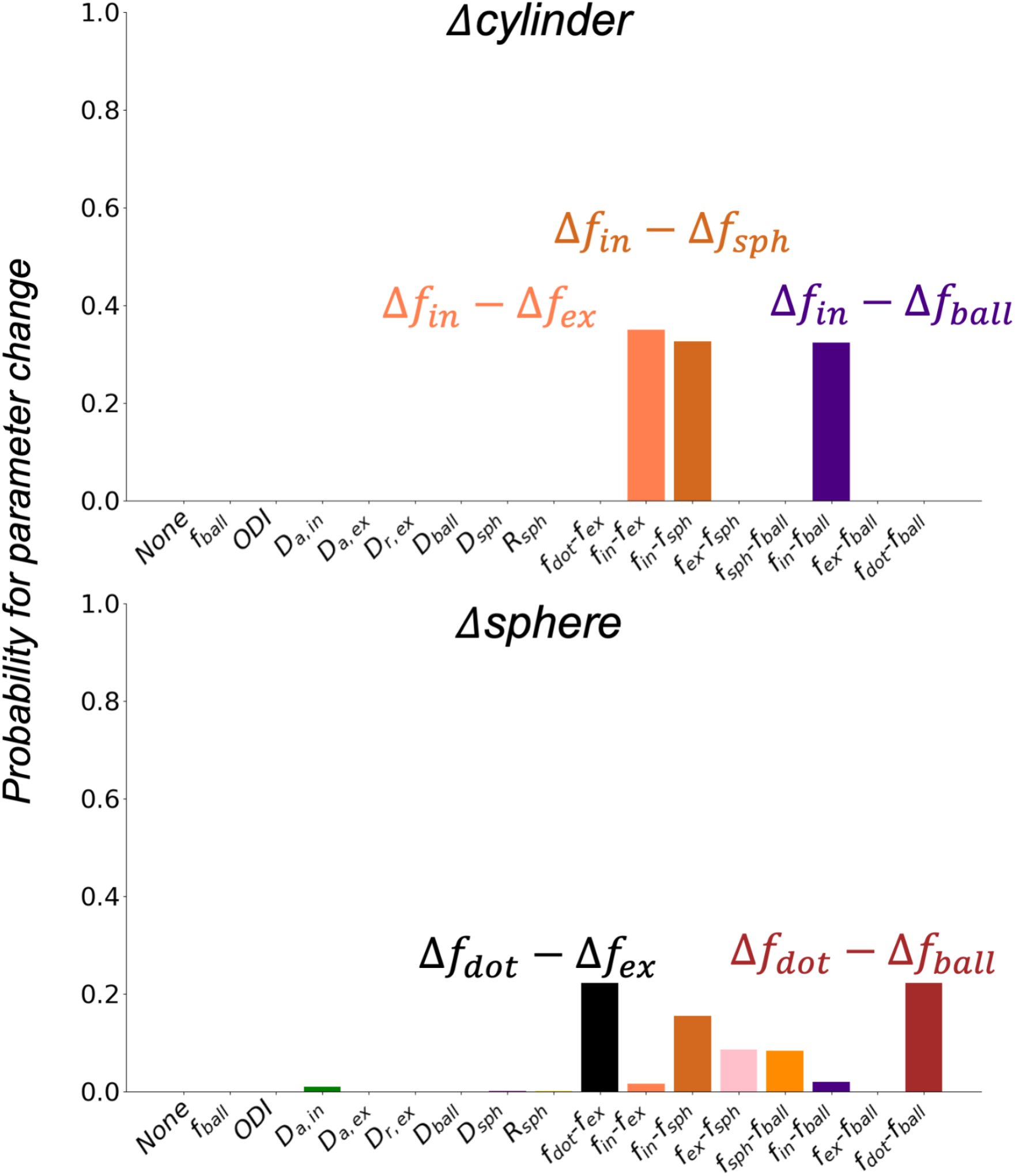
Inference of which biophysical parameter(s) best explains a change in cylinders and spheres in the mesh substrates. The change in cylinders (Δ cylinder) was best explained by the intra-axonal signal fraction replacing extra-axonal, sphere and ball signal fractions (Δ *f*_*in*_ − Δ *f*_*ex*_, Δ *f*_*in*_ − Δ *f*_*sph*_, Δ *f*_*in*_ − Δ *f*_*ball*_). The change in spheres (Δ sphere) was attributed to an increase in the dot signal fraction replacing the extra-axonal and ball signal fraction (Δ *f*_*dot*_ − Δ *f*_*ex*_, Δ *f*_*dot*_ − Δ *f*_*ball*_), though with lower probabilities relative to Δ cylinder.

### 4.3 The GLM

The GLM (c.f. Section 2.3) was set up with the microscopy metrics as the explanatory variables. For simulated data, the explanatory variables were the number of spheres and cylinders in each substrate. For the real (ex-vivo) data, the explanatory variables were the four microscopy-derived metrics: the Nissl SAF, Nissl ODI, myelin SAF, and myelin ODI.

The GLM outputs the baseline dMRI signal ***β***_***mean***_ and the direction of change in different metrics, for both the simulated data (***β***_***cylinders***_, ***β***_***spheres***_) and the ex-vivo data (***β***_***NisslSAF***_, ***β***_***NisslODI***_, ***β***_***myelinSAF***_, ***β***_***myelinODI***_). For ex-vivo data, we first investigated how each microscopy-derived metric individually explained variance in the dMRI data by performing a separate GLM for each metric (termed “single regressor”). We then determined how each metric uniquely explains variance in the dMRI data by performing multiple linear regression with all metrics in the same GLM design matrix (termed “all regressors”).

As BENCH requires the measured change in the dMRI signal **Δy** rather than ***β***_***l***_ (where 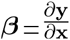), ***β***_***l***_ was multiplied with the effect size *δ*_*l*_. In the simulated data, *δ*_*l*_ was equal to the range of cylinders (n=57) and spheres (n= 328), respectively. For the ex-vivo data, the value of *δ*_*l*_ was calculated by taking the difference between the 99.9th percentile and the 0.1th percentile of each microscopy-derived metric’s distribution across the brain.

For the simulated data, the noise covariance ***σ***_***l***_ was taken to be the covariance of ***β***_***l***_ computed from 100 different instances of additive Rician noise (SNR=150). In the ex-vivo data, the noise covariance was derived from the data via a bootstrapping method (2000 iterations). In each iteration, we sampled 1000 voxels from the white matter mask and computed ***β***_***NisslSAF***_, ***β***_***NisslODI***_, ***β***_***myelinSAF***_ and ***β***_***myelinODI***_. ***σ***_***l***_ was taken to be the covariance of ***β***_***l***_ across iterations.

### 4.4 Inference

The measured change in dMRI signal due to different continuous variables (***β***_***l***_*δ*_*l*_), along with their noise covariance term (***σ***_***l***_), were used for inference. In the simulated data, we present the inferred probability averaged across 100 iterations. For ex-vivo data, the final probabilities were generated from a single inference.

## 5 Results

### 5.1 Numerical simulations

Continuous BENCH was applied to simulated data from mesh substrates with differing numbers of cylinders and spheres (Figure 5) to validate its ability to attribute these changes to appropriate biophysical model parameters (i.e. different *f*_*in*_ and *f*_*sph*_ or *f*_*dot*_, where *f*_*sph*_ and *f*_*dot*_ may be indiscernible at long diffusion times). For simulations with increasing numbers of cylinders, BENCH predicted increasing intra-axonal signal fractions replacing sphere, extra-axonal and ball signal fractions, specifically either (*f*_*in*_ − *f*_*sph*_), (*f*_*in*_ − *f*_*ex*_), or (*f*_*in*_ − *f*_*ball*_), all with probabilities ∼ 33%. For simulations with increasing numbers of spheres, continuous BENCH mainly inferred changes in the dot compartment, specifically (*f*_*dot*_ − *f*_*ex*_) and (*f*_*dot*_ − *f*_*iso*_), with probabilities ∼25%.

### 5.2 Ex-vivo macaque brain data

Continuous BENCH was applied to ex-vivo macaque data to relate changes in biophysical model parameters to myelin and Nissl SAF and ODI derived from microscopy. We explored both each metric independently (“single regressor”) and all metrics together (“all regressors”).

#### Single regressor

Figure 6 shows the inference of each parameter in separate GLMs. Note that none of the inferences have been thresholded—the plots display the inferred probabilities as they are. An increase in myelin SAF was linked to an increase in intra-axonal signal fraction replacing the extra-axonal signal fraction (Δ *f*_*in*_ − Δ *f*_*ex*_) (probability ∼ 100%). The increased density of Nissl was linked to the intra-axonal signal fraction being replaced by the sphere compartment signal fraction (Δ *f*_*sph*_ − Δ *f*_*in*_) (probability ∼ 100%). An increase in either myelin- or Nissl-derived dispersion (ODI) led to an increase in the extra-axonal signal fraction and a complementary decrease in the intra-axonal signal fraction (Δ *f*_*ex*_ − Δ *f*_*in*_) (probability ∼ 100%).

**Figure 6:**
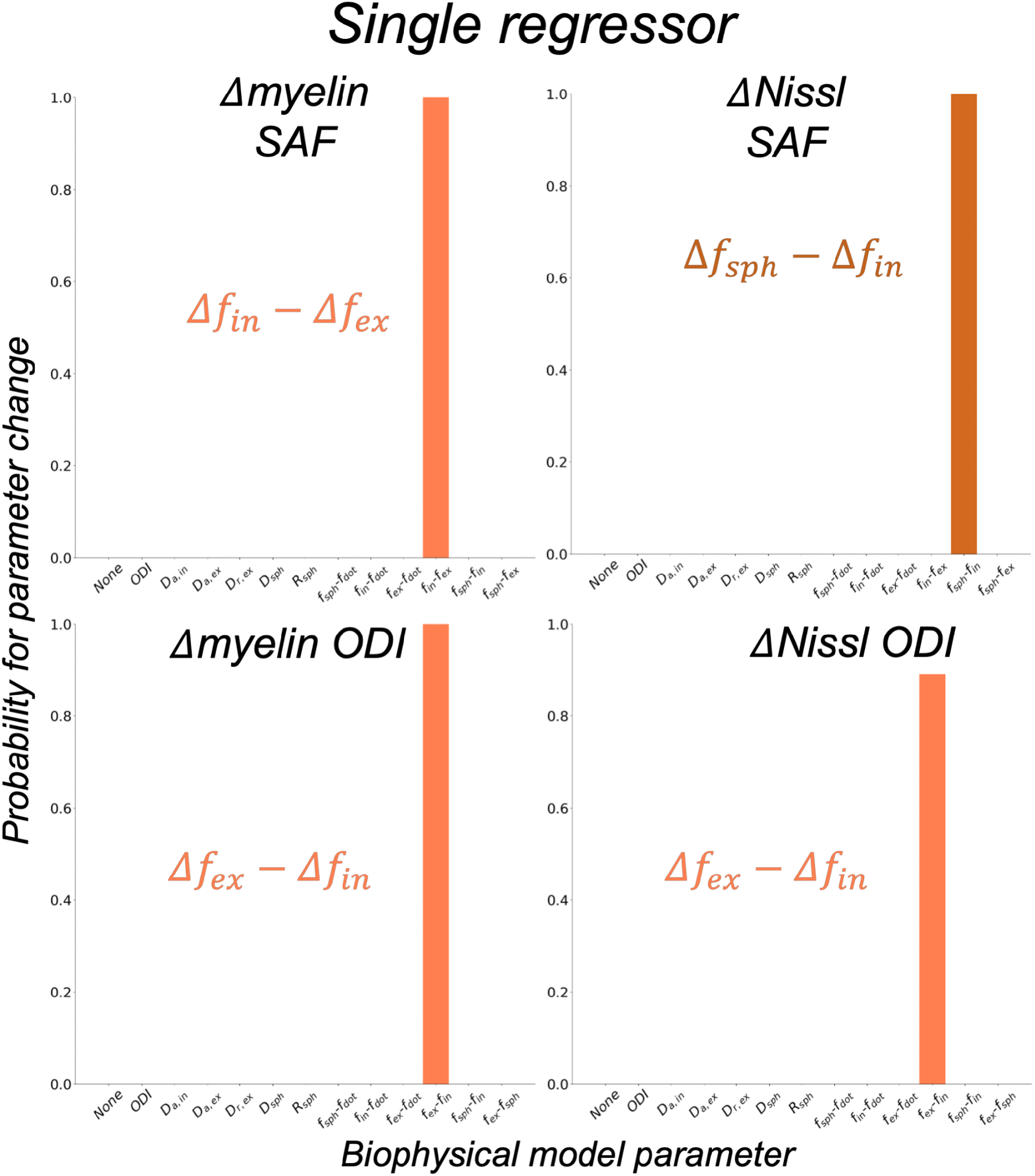
Inferred probabilities on real data when using each microscopy-derived metric (one plot per metric) as the only regressor in the GLM. Nissl in the white matter primarily represents glial cell soma. The change in myelin and Nissl density (SAF; stained area fraction) was linked to an increase in *f*_*in*_ and *f*_*sph*_ respectively. A change in either myelin or Nissl orientation dispersion index (ODI) was attributed to an increase in *f*_*ex*_. In the bottom two plots, the “Δ *f*_*in*_ − Δ *f*_*ex*_ “label on the x-axis was replaced with” Δ *f*_*ex*_ − Δ *f*_*in*_ “to reflect that the observed signal change was in the opposite direction.

#### All regressors

When using all metrics simultaneously as regressors in the GLM, the inferred probabilities for myelin-derived ODI and Nissl SAF differed (Figure 7) from the analysis using each metric independently. Myelin-derived ODI was confidently linked to model parameter ODI (probability ∼95%), while an increase in Nissl SAF was associated with increased extra-axonal signal fraction replacing the sphere signal fraction (Δ *f*_*ex*_ − Δ *f*_*sph*_) (probability ∼100%).

**Figure 7:**
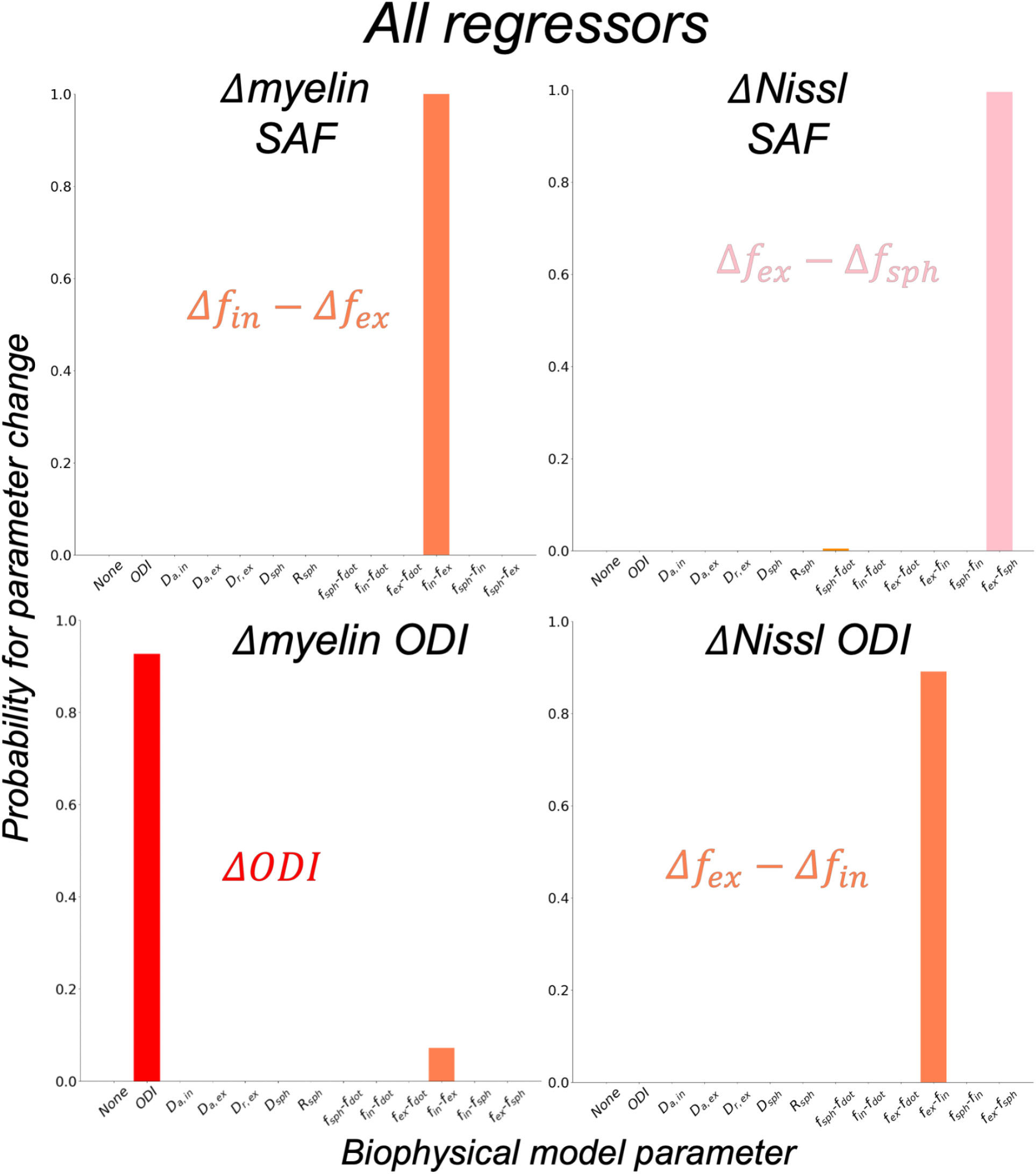
Inferred probabilities when using all microscopy-derived metrics as regressors in the same GLM. Results are shown as similarly described in Figure 6.

We further investigated these results by plotting the change in each dMRI summary measure with respect to the microscopy (Figure 8, left, titled “Measured”). This observed “pattern of change” is compared to BENCH’s trained change models showing the predicted pattern of change for each biophysical parameter (Figure 8, right, titled “Predicted”). When the measured pattern of change matches one of the predicted patterns of change, BENCH should infer that biophysical parameter with high probability. This is observed in the myelin SAF and ODI, where the measured patterns are highly similar to those from Δ *f*_*in*_ − Δ *f*_*ex*_ and ODI, respectively. Similarly, Nissl ODI was most similar to the inverted pattern of change for Δ *f*_*in*_ − Δ *f*_*ex*_, indicating a change in Δ *f*_*ex*_ − Δ *f*_*in*_. The patterns of change for myelin SAF and Nissl ODI are very similar but opposite in magnitude, which is why they were matched to Δ *f*_*in*_ − Δ *f*_*ex*_ and Δ *f*_*ex*_ − Δ *f*_*in*_ (the inverse). Though the myelin ODI also looks similar, it differs in that the changes in the *b*_7_ and *b*_10_ mean terms (1st and 3rd columns) are effectively zero, whereas the myelin SAF and Nissl ODI show small but non-zero changes in these terms. This difference (i.e. the lack of change in the *b*_7_ and *b*_10_ mean terms) results in myelin ODI matching with the Δ ODI biophysical parameter. This is as expected as changes in signal fractions, but not dispersion, would affect the *b*_7_ and *b*_10_ mean terms (equivalent to the spherically averaged signal), whilst dispersion affects the *l*2 terms only.

**Figure 8:**
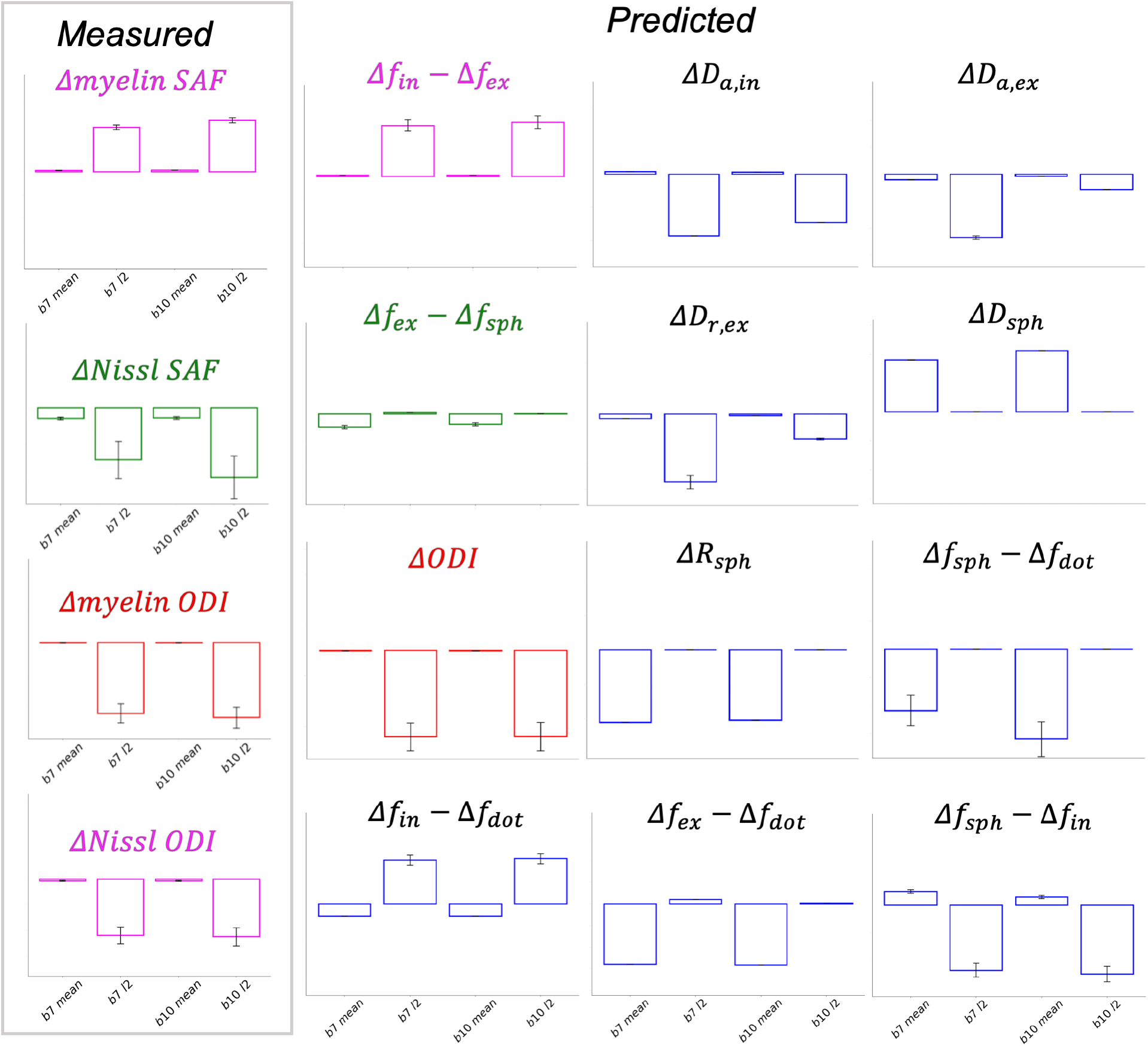
The “pattern of change” in dMRI summary measures (x-axis) for each microscopy metric from the real ex-vivo data (left box) are compared to those predicted from BENCH’s trained change models. In each plot, the dMRI summary measures on the *x* -axis are the *b*_7_ mean, the *b*_7_ *l*2, the *b*_10_ mean and the *b*_10_ *l*2 terms. The *y* -axis describes the change in signal for each summary measure with respect to a unit change in the continuous variable (left “Measured”) or a unit change in the model parameter (right, “Predicted”). The values along the *y*-axis are omitted as BENCH only considers the relative pattern of change across the summary measures, not the absolute magnitude. The colours (pink, green and red) show which patterns of change are inferred by BENCH to be most similar. Note, the myelin SAF is linked to (Δ *f*_*in*_ − Δ *f*_*ex*_), while Nissl-derived ODI is linked to same pattern of change, but in the opposite direction (i.e. Δ *f*_*ex*_ − Δ *f*_*in*_).

The Nissl SAF was more challenging to interpret, as the pattern of change did not match any of the predicted patterns of change. BENCH assigned Δ *f*_*ex*_ − Δ *f*_*sph*_ as the most similar match. Here, BENCH likely matched them based on the similarity of the *b*_7_ and *b*10 mean. The *l*2 terms look quite different. As there is high standard deviation in Nissl SAF’s *b*7 and *b*10 *l*2 summary measures, BENCH will downweight the importance of matching these terms. Ultimately, this result suggests our biophysical model is inadequate at explaining Nissl variation across the brain. This result also highlights the importance of interpreting the inferred parameters output from BENCH alongside these patterns of change; as the output probabilities collectively sum to 1, some probabilities may be inflated. This is especially the case for inferences relating to Nissl SAF.

## 6 Discussion

In this work, we modified BENCH to relate continuous variables to biophysical parameters in highly degenerate modelsWe combine continuous BENCH with an extended dMRI standard model of white matter to investigate dMRI sensitivity to axons and glia. We first applied continuous BENCH to simulated data, where the parameters inferred by BENCH accurately reflected the simulated changes in spheres/cylinders in mesh substrates, validating our approach. We then applied continuous BENCH to real ex-vivo macaque MRI data with co-registered maps of microscopy metrics for myelin and cell bodies acquired from the same brain.

We demonstrate that myelin SAF, a proxy for myelinated fibre density, is related to *f*_*in*_, and myelin ODI related to model parameter ODI. In addition, we found evidence that glia are poorly represented by our extended standard model. In addition to demonstrating continuous BENCH as a method for probing microstructural correlates ex-vivo, we provide a first demonstration of its application to continuous variables.

### 6.1 Numerical simulations

Our results from the numerical simulations demonstrate continuous BENCH when the ground truth is known. When applied to mesh substrates with different numbers of cylinders (representing axons) or spheres (glia soma), continuous BENCH inferred the most probable change to be an increase in *f*_*in*_ and *f*_*dot*_, respectively. Cylinders are linked to a change in *f*_*in*_, accompanied by reductions in the other compartments. This may be explained by the relatively low packing densities in some substrates, where diffusing spins that encounter few membranes can be described by free (*f*_*ball*_) rather than anisotropic (*f*_*ex*_) diffusion. Increasing spheres is linked to an increase in *f*_*dot*_, at the expense of the extra-axonal (*f*_*dot*_ − *f*_*ex*_) and ball (*f*_*dot*_ − *f*_*ball*_) compartment. The dot compartment is typically used to model water trapped in very small isotropic spaces to the point where they appear stationary or with negligible diffusion. Our simulation uses a reduced bulk diffusivity (0.8 *μm*^2^/ms) for ex-vivo tissue conditions, where at long diffusion times, a sphere may be indistinguishable from a dot compartment. The dot compartment’s replacement by the extra-axonal or ball compartment may again imply the unrealistic packing densities of our substrates, as earlier suggested. These results confirm that continuous BENCH can accurately infer changes in soma- and axon-like structures, with the assumptions of impermeable membranes and high SNR (SNR=150).

### 6.2 Ex-vivo macaque brain data

When continuous BENCH was applied to real data from an ex-vivo macaque brain, the myelin SAF, a measure of myelin density, was predominantly linked to an increasing intra-axonal compartment replacing the extra-axonal compartment (*f*_*in*_ − *f*_*ex*_). This follows expectations, as myelinated fibres are thought to be well described by stick-like diffusion in this regime [8, 30].

When considered separately, BENCH linked each microscopy-derived dispersion metric to an increased extra-axonal compartment replacing the intra-axonal compartment (*f*_*ex*_ − *f*_*in*_). However, when considering all microscopy metrics simultaneously as regressors, the biophysical model parameter ODI was found to best explain the variation in myelin-derived ODI (Figures 7, 8). This confirms the model’s ODI’s specificity in mapping the dispersion of myelinated axons. The apparent relationship with the extra-axonal space (uncovered in the “single regressor” analysis) was found to be primarily driven by a strong anticorrelation between myelin SAF and ODI, and a high correlation between myelin and Nissl ODI (data not shown), highlighting the importance of accounting for the covariance in different microstructural features during MRI-microscopy comparisons.

In both regression analyses—”single regressor” and “all regressors”—Nissl-derived ODI was associated with *f*_*ex*_ − *f*_*in*_. An increase in the extra-axonal space may be associated with fibre geometries that are more dispersed and therefore less tightly packed. Hence, our results may suggest Nissl-derived ODI as an indirect measure of fibre dispersion, albeit less specific than myelin-derived ODI. Nissl orientations are derived via a structure tensor which outputs orientations similar to those from myelin stains. This similarity may arise because glial cells cluster in the spaces between axons, creating an apparent orientation that mirrors the axonal orientation [31]. However, exactly how we should interpret these orientations requires more investigation, where there may be important differences between myelin- and Nissl-derived ODI (i.e. two metrics are not equivalent). Notably here, the myelin ODI was linked to the standard model ODI parameter, whilst the Nissl ODI was not. This warns against their interpretation as being equivalent measures of microstructure dispersion,

Interpreting the Nissl SAF is more challenging. Nissl stains the cell soma of both neurons and glia indiscriminately. Since there are very few neuronal soma in WM, our Nissl SAF likely reflects glial cell soma density. Investigation of the dMRI patterns of change (Figure 8, all regressors) revealed that none of the change models’ predicted patterns of change adequately matched that from the Nissl SAF. In this case, BENCH infers the parameter with the highest probability (here *f*_*ex*_ − *f*_*sph*_), although it may not be a good fit and the probability (normalised across all parameters) may be inflated. The lack of a match suggests the extended standard model, in its current form, is incapable of describing signal variation due to glia cell soma (Nissl SAF). This could be addressed in BENCH by incorporating more complex biophysical models (e.g. with exchange or inter-compartmental variations in *T*_2_), or more complex change models (e.g. with changes in more than two microstructure compartments at the same time, or the simultaneous change of a signal fraction and ODI). Nonetheless, our results show that there is non-negligible dMRI signal sensitivity to changes in glial cell soma, after accounting for myelin load and dispersion. This finding is in agreement with recent evidence of dMRI’s sensitivity to glia [17, 32-35].

### 6.3 Limitations

There are limitations to this study. First, our numerical simulation only validates continuous BENCH’s ability to accurately infer simple changes in intra-soma and intra-axonal compartments in specific conditions, such as the case where membranes are impermeable and data have high SNR (SNR=150). These conditions may not be met when applying continuous BENCH to ex-vivo data, where some assume glia to be in fast exchange with the extra-axonal space [36, 37], though exchange is likely attributed to glial processes rather than soma [30]. Second, the mesh substrates used for simulations are not very biologically realistic. Future validation work would benefit by incorporating more realistic packing densities, cellular morphologies [38, 39] and axon diameters (i.e. ∼ 1-2 *μ*m [15]). Third, we do not incorporate other microscopy-derived metrics that may relate to dMRI. For example, we do not include metrics derived from neurofilament staining, which may better explain variation in intra-axonal signal fraction and/or ODI, as our myelin metrics may miss contributions from low/unmyelinated axons.

## Authors’ contributions

**DZLK** conceived the study design and implemented continuous BENCH, which involved adapting the code from the original BENCH framework. DZLK also conducted data simulation, data analyses, and drafted the manuscript. **HR** modified the BENCH framework and helped interpret results from data analyses. **INH** developed the TIRL registration platform and performed the MRI-microscopy co-registration. **AS** processed and curated microscopy data. **SJ** conceived the study design and contributed to the design of continuous BENCH. **MC** contributed to the design of continuous BENCH and helped interpret results from data analyses. **KLM** conceived the study design, contributed to all data analyses, and helped draft the manuscript. **AFDH** conceived the study design, contributed to implementation of continuous BENCH, contributed to all data analysis, and helped draft the manuscript.

## Data availability

The BigMac dataset is openly available via the Digital Brain Bank:

https://open.win.ox.ac.uk/DigitalBrainBank/. The code will be made available upon publication.

## Acknowledgements

AFDH and KLM contributed equally to this work. Data acquisition was funded by the grant MR/K02213X/1 from the Medical Research Council (MRC). DZLK was supported by the Hrothgar Singaporean Clarendon Scholarship and the Nuffield Department of Clinical Neurosciences studentship. AFDH and INH were supported by the EPSRC, MRC and Wellcome Trust (grants EP/L016052/1 and WT202788/Z/16/A). INH was also supported by the Clarendon Fund in partnership with the Chadwyck-Healey Charitable Trust. KLM, SJ and MC are supported by the Wellcome Trust (grants WT202788/Z/16/A, WT215573/Z/19/Z and WT221933/Z/20/Z). The Wellcome Centre for Integrative Neuroimaging is supported by core funding from the Wellcome Trust (203139/Z/16/Z) and (203139/A/16/Z). For the purpose of open access, the author has applied a CC BY public copyright licence to any Author Accepted Manuscript version arising from this submission.

## Appendices Summary measures for diffusion MRI data

dMRI summary measures were calculated using the spherical harmonic decomposition [20] of the dMRI signal. Here, the dMRI signal can be decomposed into a unique linear combination of orthonormal basis functions over the surface of a unit sphere (spherical harmonics):

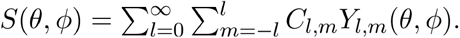

The dMRI signal *S*, parameterised by the polar (*θ*) and azimuthal (*ϕ*) angles of the standard spherical coordinate system, is now described as a linear combination of coefficients (*C*) and spherical harmonics (*Y*) up to a pre-decided degree *l* = 0, 1, 2, … and order *l* = −*l*, … *l*. Since S is symmetric around the origin, only even-degree harmonics (i.e., *l* = 0, 2, 4, …) have non-zero coefficients.

We then computed rotationally invariant measures for each shell of dMRI signal as:

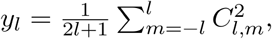

where *yl* is the summary measure at degree *l* for a single shell of data [1, 21]. Here, we set *l* = 0, 2, (i.e. 0th and 2nd degree), since a higher *l* quantifies the signal anisotropy, but at a much lower SNR. Lastly, we took the mean over the norm to ensure that the scale remains constant across degrees. With this formulation, we summarised all measured dMRI signal data points with 4 rotationally invariant summary measures (2 summary measures × 2 shells) per voxel.

### Mask for ex-vivo macaque brain data

**Figure S1:**
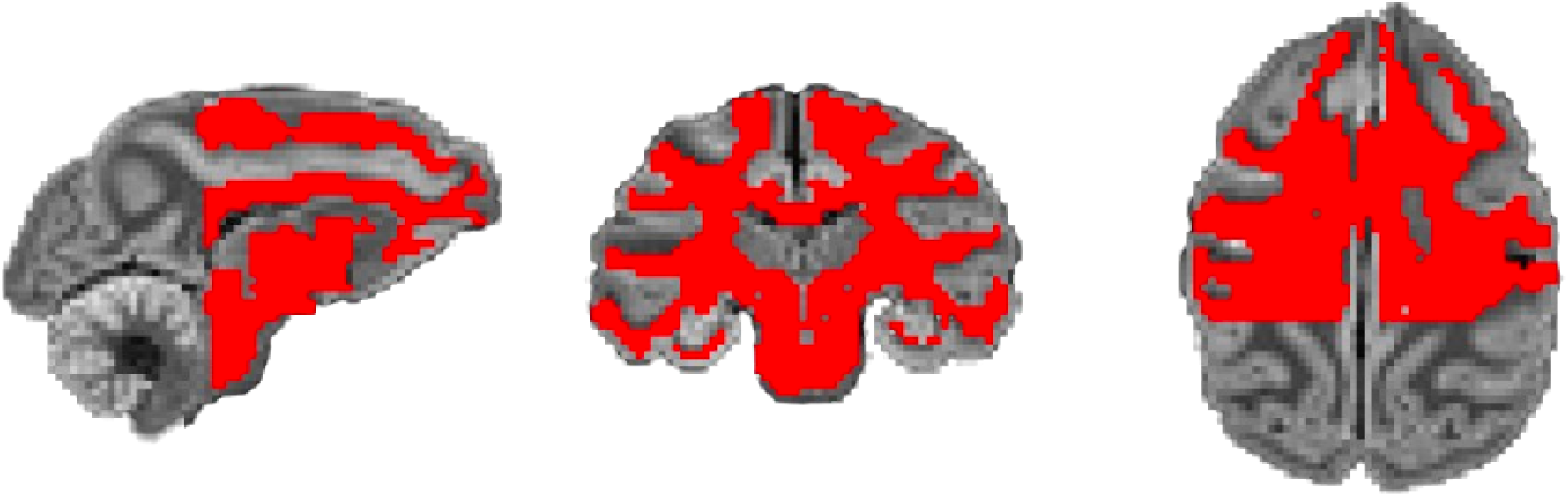
The final WM mask (red) containing the 17945 voxels used for analysis. The mask is overlaid on the ex-vivo dMRI data of the macaque brain.

